# Exploring information exchange between *Thesium chinense* and its host *Prunella vulgaris* through joint transcriptomic and metabolomic analysis

**DOI:** 10.1101/2023.12.27.573399

**Authors:** Anping Ding, Zengxu Xiang, Ruifeng Wang, Juan Liu, Wenna Meng, Yu Zhang, Guihong Chen, Gang Hu, Mingpu Tan

**Affiliations:** College of Horticulture, Nanjing Agricultural University, Nanjing, Jiangsu, China; College of Life Sciences, Nanjing Agricultural University, Nanjing, Jiangsu, China

**Keywords:** *T. chinense*, *P. vulgaris*, transferred metabolite, mobile gene, haustoria formation

## Abstract

Background: *Thesium chinense* known as “plant antibiotic” is a facultative root hemi-parasitic herb while *Prunella vulgaris* can serve as its host. However, the molecular mechanisms underlying the communication between *T. chinense* and its host remained largely unexplored. The aim of this study was to provide a comprehensive view of transferred metabolites and mobile mRNAs between *T. chinense* and *P. vulgaris*. Results: The wide-target metabolomic and transcriptomic analysis identified 5 transferred metabolites and 668 mobile genes between *T. chinense* and *P. vulgaris*, as well as haustoria formation related 56 metabolites and 189 genes. Furthermore, we inferred a regulatory network that might be involved in haustoria formation, and 18 genes promoting haustoria formation and 1 gene inhibiting it were identified as a consequence. There were 4 metabolites (ethylsalicylate, eriodictyol-7-O-glucoside, aromadendrin-7-O-glucoside and pruvuloside B) that are transferred from *P. vulgaris* to *T. chinense*, whereas 2-ethylpyrazine was transferred from *T. chinense* to *P. vulgaris*. Conclusions: These results suggested that there was an extensive exchange of information with *P. vulgaris* including transferred metabolites and mobile mRNAs, which might facilitate the haustoria formation and parasition of *T. chinense*.

**Author summary:** *T. chinense* known as “plant antibiotic” is a facultative root hemi-parasitic herb while *P. vulgaris* can serve as its host. Currently, the information exchange between *T. chinense* and its host remained unknown, and a comprehensive view of transferred metabolites and mobile mRNAs between *T. chinense* and its host is critical so that appropriate chemical and genetic improvement can be used to facilitate haustoria formation and successful parasitism. Here, we employ the conjoint wide-target metabolomic and transcriptomic analysis to explore the information exchange between *T. chinense* and *P. vulgaris.* We identified 5 transferred metabolites and 668 mobile genes between *T. chinense* and *P. vulgaris*, as well as haustoria formation related 56 metabolites and 189 genes. Our study provides new insights into the complex interplay between parasite and host during parasitism.

## Introduction

Parasitic plants are greatly diverse, which are currently categorized on two main bases: holoparasite and hemiparasite according to whether they are able to photosynthesis or not, and rootparasite and stemparasite according to the location of the parasitism on the host plant ^[1,2]^. Despite their remarkable diversity, all parasitic plants share a unique specialized organ called the haustorium ^[3]^, which was described as “the essence of parasitism” ^[4]^. During the interaction with their hosts, the haustorium exhibits dynamic changes in its functions. Early in the commensal process, the haustorium aids parasite in host attachment and invasion, and subsequently, it facilitates the uptake of nutrients, hormones and signaling molecules ^[5]^. The resulting symplastic continuity enables macromolecules or genetic materials such as RNA to be transferred between the hosts and the parasites ^[6]^.

*T. chinense* is a hemiparasitic plant of the genus *Thesium* in the Santalaceae family, which is widely distributed in China, Japan, and Korea ^[7]^. Modern pharmacological researches have proved that *T. chinense* has diverse activities including anti-inflammation ^[8,9]^, antimicrobial effect ^[10–13]^, analgesic activity ^[14]^, antioxidant activity ^[15]^ and anti-nephropathy ^[16]^. Moreover, *T. chinense* known as “plant antibiotic” ^[17]^ has a great deal of efficacy for the treatment of mastitis, tonsillitis, pharyngitis, pneumonia, and upper respiratory tract infections ^[11,18,19]^. The hosts of *T. chinense* are distributed extensively in many plant families ^[20]^, including *P. vulgaris*, a perennial herb in the family Lamiaceae ^[21]^. *T. chinense* is a root-parasitic plant, which is attached to the roots of *P. vulgaris* by the haustoria to sustain its own growth and development.

Due to their unique ways of symbiosis, the parasitic plants not only absort water ^[22]^ and nutrients from the host, but also leverage secondary metabolites, mRNA ^[23,24]^, proteins ^[24]^, and systemic signals ^[25,26]^ from their hosts. *Cistanche deserticola* utilizes its host *Haloxylon ammodendron* derived metabolites for better survival ^[27]^. *Cuscuta* not only transmits mRNAs between different host plants ^[28]^, but also exchanges proteins with its hosts, and even proteins with the same functions can be transferred between different host plants of *Cuscuta* ^[29]^. Parasites plant may also actively transfer phytohormones to the hosts to manipulate host physiology ^[25]^. In recent years, the researches of *T. chinense* focused on the regulation of seed dormancy-breaking growth and development ^[30]^, in vitro anti-inflammatory and antimicrobial activity of extracts ^[8,11,31,32]^, host range and selectivity ^[33]^, and developmental reprogramming of haustoria formation ^[34]^. In contrast, there are fewer studies on the exchange of information between *T. chinense* and host plants. Therefore, it is necessary to further investigate the molecular mechanism governing the interaction crucial for successful parasitism and the subsequent symbiosis between parasitic plants and the host.

To explore the information exchange events between *T. chinense* and *P. vulgaris*, we conducted an integrated wide-target metabolomic and transcriptomic analysis, especially during the established parasitism relationship between *T. chinense* and its host, *P. vulgaris*. In this sutdy, 5 transferred metabolites and 668 mobile genes were identified between *T. chinense* and *P. vulgaris*, as well as haustoria formation related 56 metabolites and 189 genes. By comparing the gene expression profiles and metabolic profiles of both partners, we aim to identify key genes and metabolites involved in the establishment and maintenance of the parasitic relationship. This study not only explored the information exchange events between *T. chinense* and its host, *P. vulgaris*, but also discussed major understandings of haustoria formation and host invasion, shedding light on the complex interplay between parasite and host during parasitism.

## Results

### Root morphology of *T. chinense* and its host *P. vulgaris* post parasition

The root morphology of individual *T. chinense* and its host *P. vulgaris*, and the chimeric root post symbiosis were observed histologically (Fig. 1A). The result showed that there were a large number of ivory spherical haustoria at the root of *T. chinense* (Fig. 1B). Although the roots of *T. chinense* were tightly attached to the roots of *P. vulgaris*, the haustoria did not completely penetrate the roots of *P. vulgaris* (Fig. 1C), implying that the bridge between *P. vulgaris* chimera and the haustorium was undergoing changes and transmitting cargos (Fig. 1D). To explore the information exchange events between *T. chinense* and its host *P. vulgaris*, *T. chinense* chimera (THC) and *P. vulgaris* chimera (PC) from the symbiont roots, and the root counterparts of independent *T. chinense* (TH) and *P. vulgaris* (P) seedlings were collected for the subsequent transcriptomic and metabolomic analysis.

**Fig. 1.**
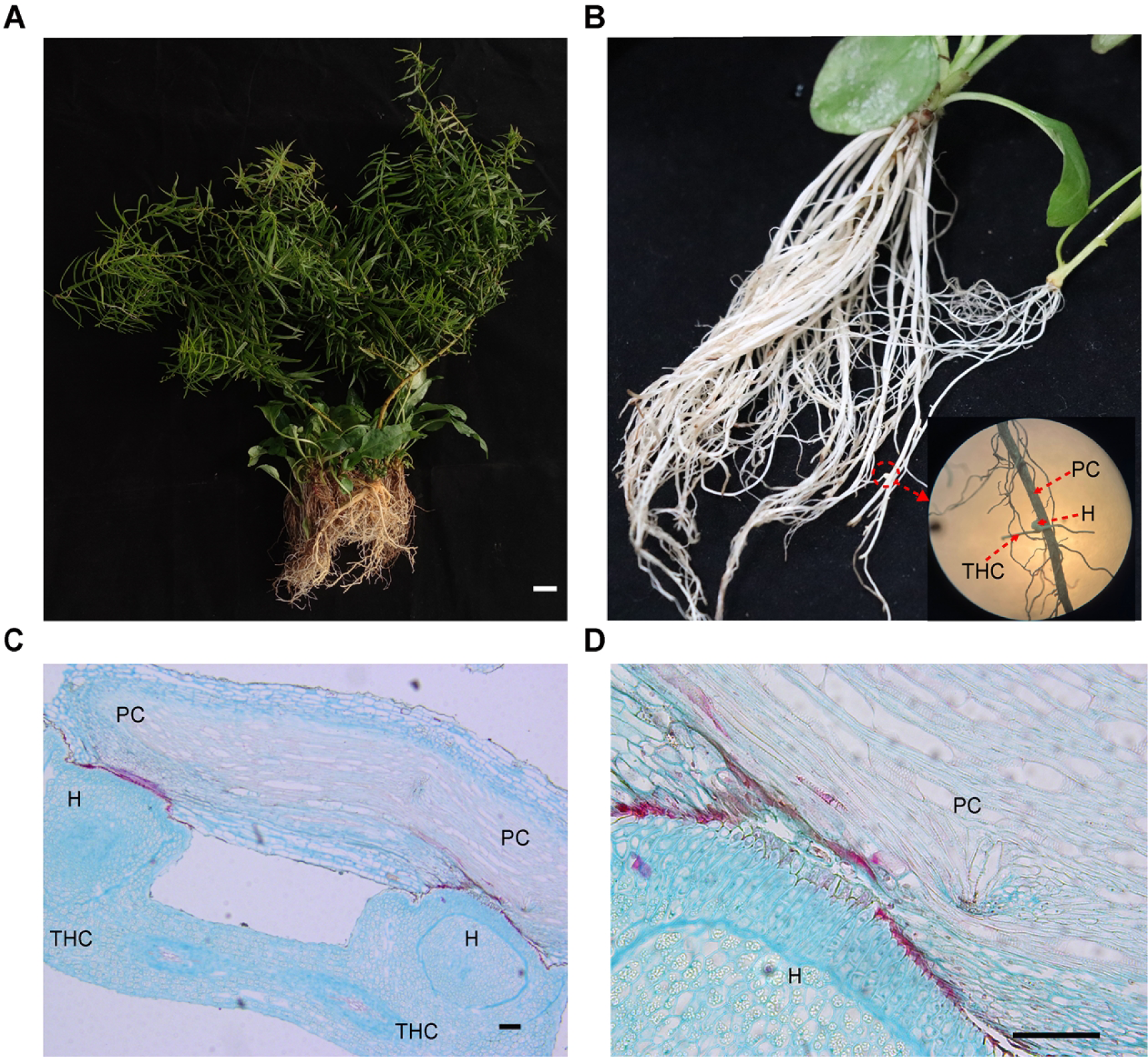
Morphology of *T. chinense* and its host *P. vulgaris* chimeric root. A: *T. chinense* and its host *P. vulgaris*. B: *T. chinense* chimera is connected to its host *P. vulgaris* chimera through haustoria. C-D: Structure of *T. chinense* chimera, haustoria and *P. vulgaris* chimera. THC: *T. chinense* chimera. H: Haustorium. PC: *P. vulgaris* chimera.

### Metabolomic changes in *T. chinense* and its host *P. vulgaris* post symbiosis

To identify the metabolites transferred between *T. chinense* and its host *P. vulgaris*, the wide-target metabolomic analysis was conducted. Consequently, 1,014 metabolites were identified in *T. chinense*, *P. vulgaris* and their chimeras (Fig. 2A and S1 Table), and the PCA analysis of these metabolites showed that they can be clearly separated into four clusters corresponding to the four sampling groups (Fig. 2B). These results suggested significantly different pattern of metabolites accumulation among TH, THC, PC, and P, emphasizing the changes caused by parasites.

**Fig. 2.**
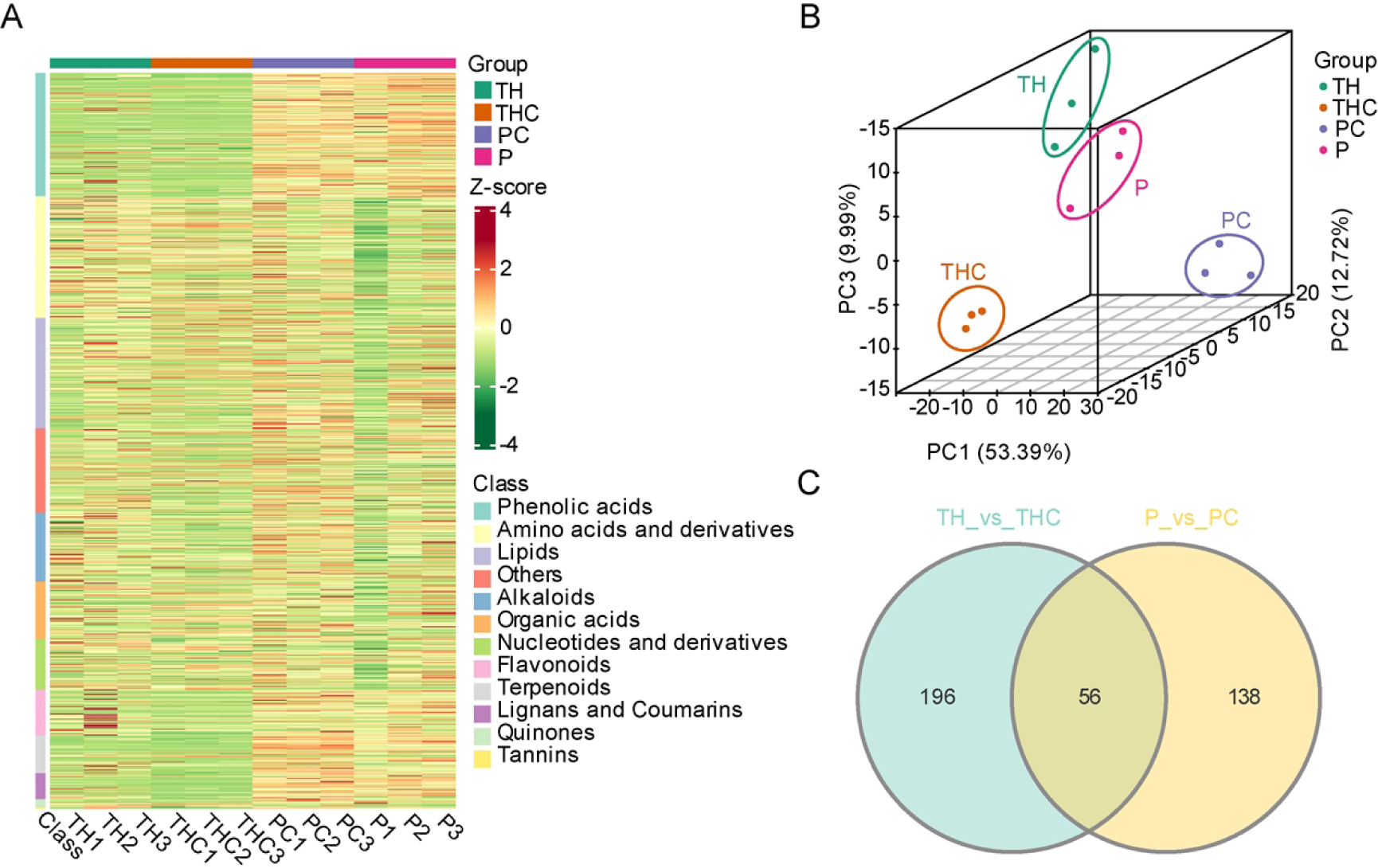
The metabolomic analysis of *T. chinense* and its host *P. vulgaris* post symbiosis. A: Heat map visualization of metabolites in *T. chinense*, *P. vulgaris* roots and their chimera. B: PCA analysis of metabolites in *T. chinense*, *P. vulgaris* roots and their chimera. C: Venn diagrams revealing the relationship of differentially accumulated metabolites (DAMs) in *T. chinense* chimera and its host *P. vulgaris* chimera. TH: *T. chinense*. THC: *T. chinense* chimera. PC: *P. vulgaris* chimera. P: *P. vulgaris*.

Then the differentially accumulated metabolites (DAMs) in *T. chinense* and its host *P. vulgaris* post symbiosis, were identified using the screening criteria of |log_2_FoldChange| ≥ 1 and VIP ≥ 1. Compared with the roots of intact *T. chinense*, 252 DAMs were identified in *T. chinense* chimera, of which 75 were upregulated and 177 were downregulated (S2 Table). Compared with the roots of parasition-free *P. vulgaris*, a total of 194 DAMs were identified in *P. vulgaris* chimera, of which 159 were upregulated while 35 were downregulated (S3 Table). Moreover, 56 common DAMs were altered both in PC and THC compared to the roots of *T. chinense* and *P. vulgaris*, therefore, these 56 DAMs can be regarded as haustoria formation related metabolites (Table 1 and Fig. 2C).

**Table 1.**
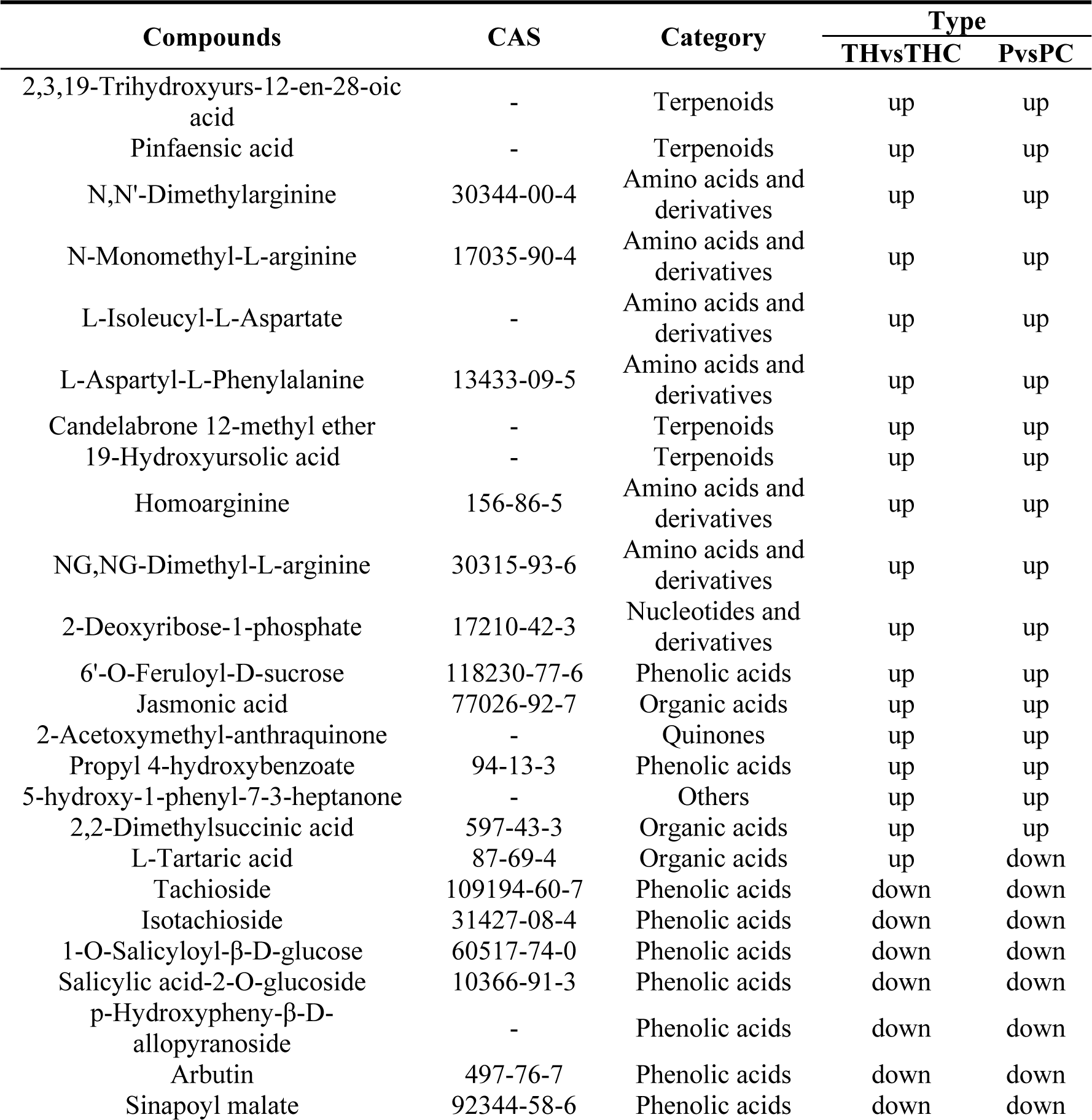
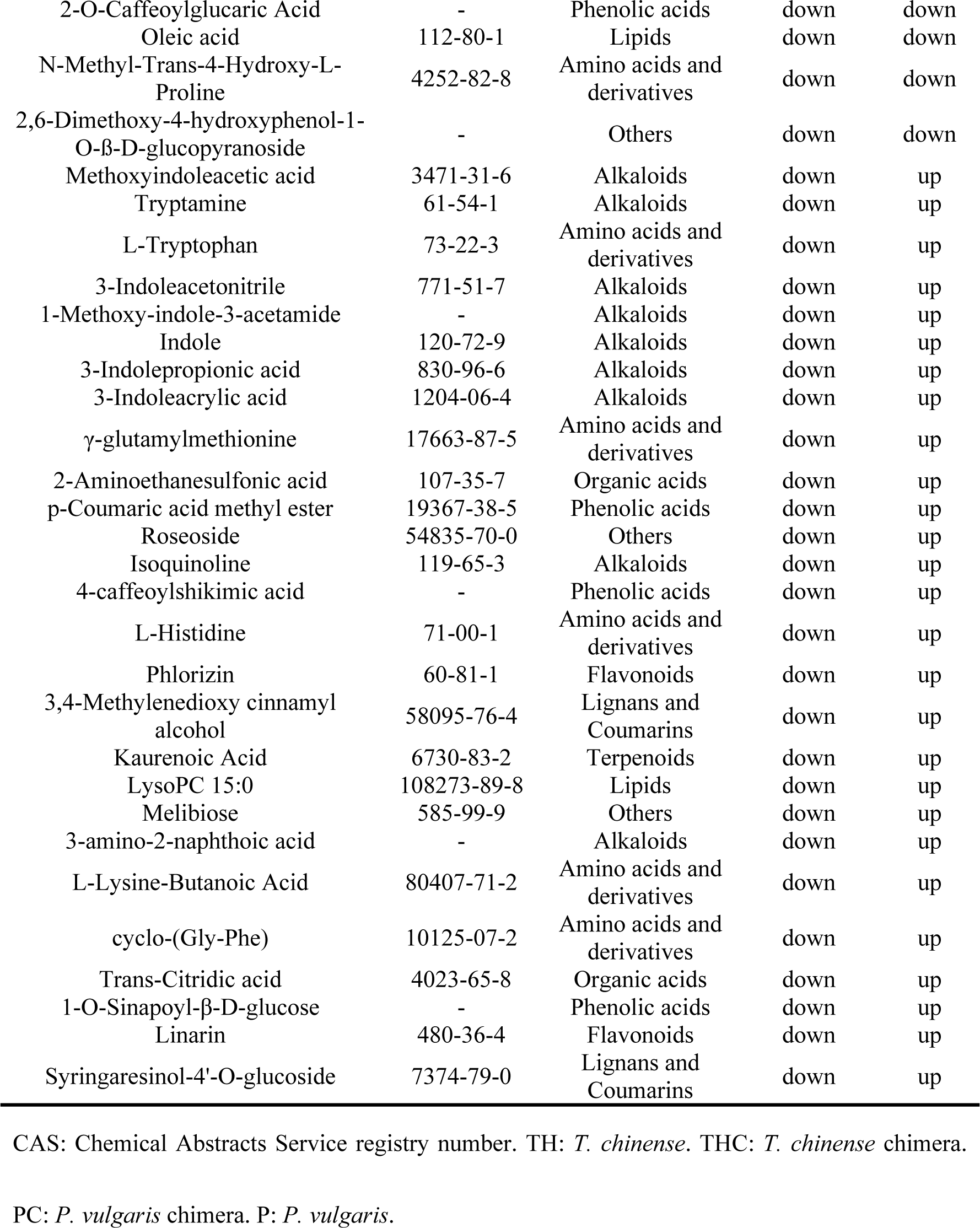
Differentially accumulated metabolites (DAMs) related to haustoria formation.

Regarding the DAMs category, phenolic acids, amino acids and derivatives, flavonoids and alkaloids accounted for more than half of the total DAMs in the TH vs THC group. Phenolic acids had the highest percentage of 27.38% (Fig. 3A), and most of the phenolic acids in *T. chinense* chimera showed a decreasing trend compared to *T. chinense*, but ethylsalicylate was upregulated in *T. chinense* chimera. A total of 23 flavonoids were detected, and most of DAMs associated with flavonoid biosynthesis including kaempferol derivatives were downregulated in *T. chinense* chimera. Another notable issue is that most of the auxin biosynthesis related components, including indole, 3-indolepropionic acid, and 3-indoleacrylic acid were downregulated in *T. chinense* chimera (S2 Table).

**Fig. 3.**
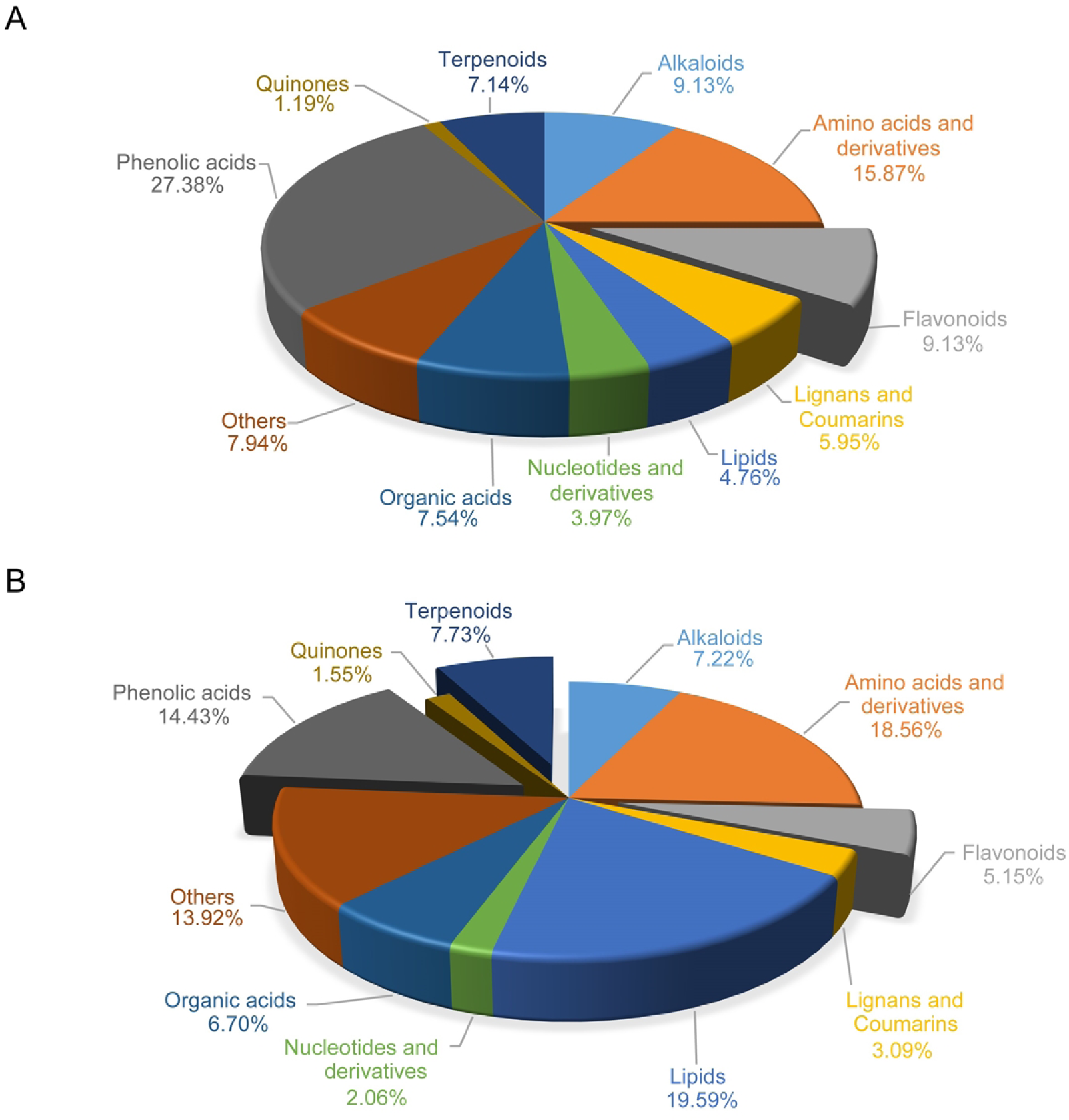
The proportion of differentially accumulated metabolites (DAMs) category in *T. chinense* chimera and its host *P. vulgaris* chimera. A: TH vs THC. B: P vs PC. TH: *T. chinense*. THC: *T. chinense* chimera. PC: *P. vulgaris* chimera. P: *P. vulgaris*.

In the P vs PC comparison group, DAMs in the category of phenolic acids, flavonoids and terpenoids accounted for 14.43%, 5.15% and 7.73%, respectively (Fig. 3B). Compared to *P. vulgaris*, ferulic acid methyl ester and p-coumaric acid methyl ester accumulated more in the *P. vulgaris* chimera, whereas the accumulation of protocatechuic acid, salicylic acid-2-O-glucoside, and arbutin was in the contrast trend. After symbiosis, the content of most flavonoids increased in *P. vulgaris* chimera. Interestingly, terpenoids (including kaurenoic acid, 18-oxoferruginol, and serratagenic acid) and jasmonic acid (JA) were all upregulated in *P. vulgaris* chimera (S3 Table).

### The exchanges of metabolites between *T. chinense* and its host *P. vulgaris* during parasitism

To explore the information exchange between *T. chinense* and its host *P. vulgaris*, the accumulation pattern of metabolites in the four groups (TH, THC, PC, and P) was compared. Consequently, those metabolites not detected in TH or P but accumulated in other three samples were defined as transferred metabolites. As a result, 5 transferred metabolites (ethylsalicylate, eriodictyol-7-O-glucoside, aromadendrin-7-O-glucoside, pruvuloside B, 2-ethylpyrazine) were identified (Table 2). Particularly pruvuloside B, a characteristic component of *P. vulgaris*, was detected in PC, P and THC, however it was not found in TH roots, suggesting a transfer of this metabolite from *P. vulgaris* chimera to *T. chinense* chimera (host→parasite direction). Similar host→parasite mobile metabolites include ethylsalicylate, eriodictyol-7-O-glucoside, and aromadendrin-7-O-glucoside. By Contrast, 2-ethylpyrazine was presented in TH, THC, and PC but not in P roots, indicating that this metabolite was transferred from *T. chinense* chimera to *P. vulgaris* chimera (parasite→host direction).

**Table 2.** The transferred metabolites between *T. chinense* and its host *P. vulgaris*.

| Compounds | CAS | Category | TH | THC | PC | P |
| --- | --- | --- | --- | --- | --- | --- |
| Ethylsalicylate | 118-61-6 | Phenolic acids | - | 35397 | 500949 | 542212 |
| Eriodictyol-7-O-glucoside | 38965-51-4 | Flavonoids | - | 29411 | 3851867 | 4694065 |
| Aromadendrin-7-O-glucoside | 28189-90-4 | Flavonoids | - | 113418 | 1191540 | 604233 |
| Pruvuloside B | - | Terpenoids | - | 1789 | 54923 | 40759 |
| 2-Ethylpyrazine | 13925-00-3 | Alkaloids | 49686 | 43058 | 78849 | - |

CAS: Chemical Abstracts Service registry number. TH: *T. chinense*. THC: *T. chinense* chimera.
PC: *P. vulgaris* chimera. P: *P. vulgaris*.

Regarding the haustoria formation related hormones, the auxin biosynthesis related components accumulated significantly more in PC, whereas the opposite was true in THC. Jasmonic acid (JA) was upregulated in both THC and PC. Moreover, another 16 metabolites were also synchronized upregulated in both chimeras groups, and these metabolites can be considered as promoting haustoria formation. Conversely, 11 metabolites downregulated in both chimeras groups may inhibit haustoria formation (Table 1 and S4 Table).

### Transcriptomic changes in *T. chinense* and its host *P. vulgaris* post symbiosis

Besides the metabolomic fluctuation, the parasitism of *T. chinense* also caused significant transcriptomic changes. To address this issue, transcriptomic profiling of the root samples of TH, THC, PC and P were conducted. Then a stringent cutoff (|log_2_FoldChange| ≥ 1 with the adjusted p-value padj < 0.05) was used to identify differentially expressed genes (DEGs) in *T. chinense*, *P. vulgaris* and their chimeras post parasition. Consequently, 11640 and 8705 DEGs were identified in the comparison of TH vs THC (S5 Table) and P vs PC (S6 Table), respectively.

To infer the biological functions of DEGs of *T. chinense* and its host *P. vulgaris* post symbiosis, the GO and KEGG enrichment analysis of DEGs were performed. Regarding the DEGs in TH vs THC group, the GO entries and proportions with the most significant enrichment in biological process, cellular component, and molecular function were photosynthesis/light reaction, photosystem and hydrolase activity/hydrolyzing N-glycosyl compounds, respectively (Fig. 4A), while the three most significant counterparts in P vs PC group were amino acid transport, ER body, and organic acid binding (Fig. 4B). The KEGG enriched pathways of DEGs in TH vs THC and P vs PC were similar, both including phenylpropanoid biosynthesis, flavonoid biosynthesis and plant hormone signal transduction. In addition, the DEGs in the TH vs THC group were also highly enriched in fructose and mannose metabolism, vitamin B6 metabolism and photosynthesis-antenna proteins (Fig. 4C). However, the highly represented pathways of DEGs in P vs PC were plant-pathogen interaction and MAPK signaling pathway-plant (Fig. 4D).

**Fig. 4.**
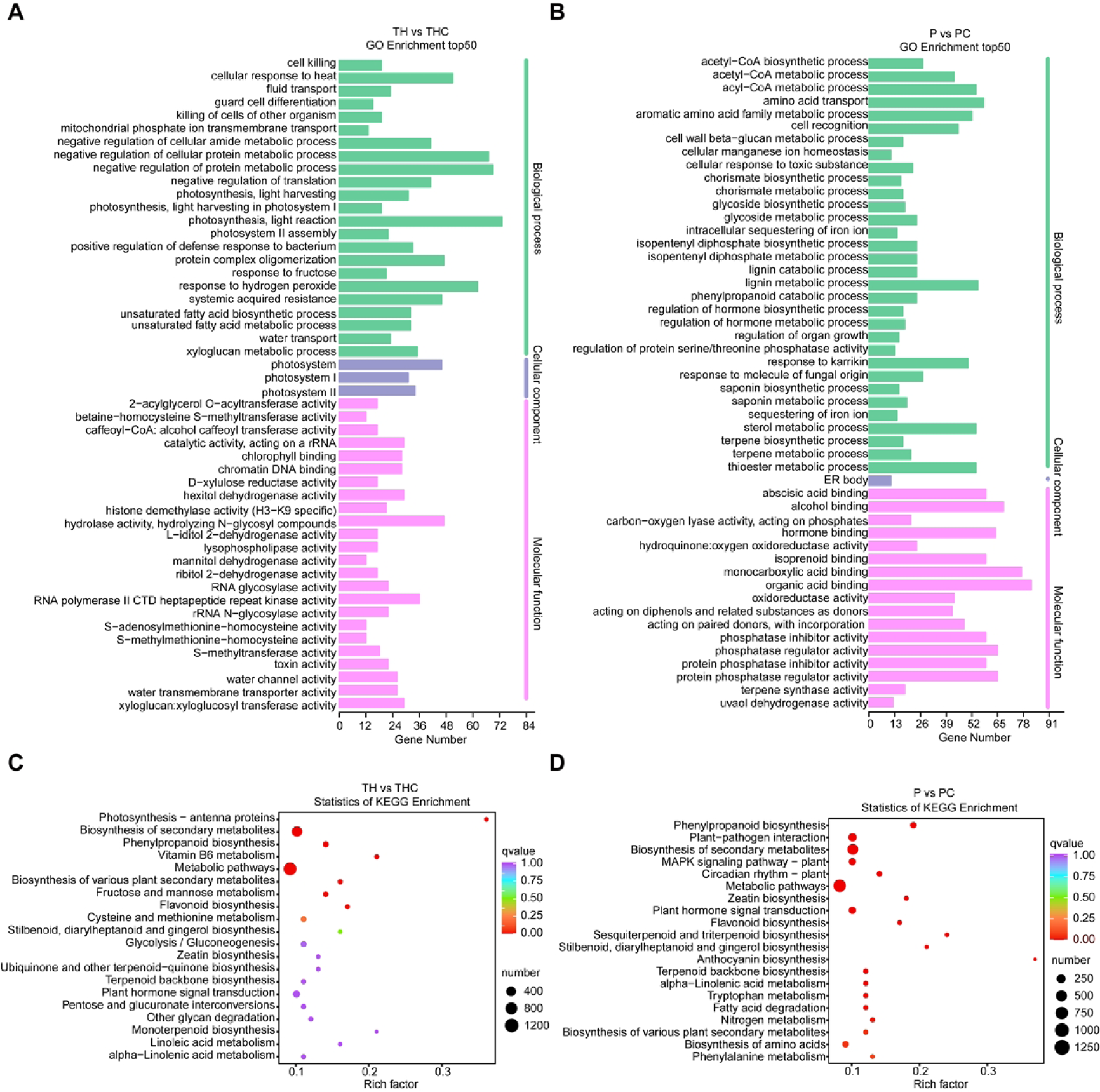
The enrichment analysis of DEGs based on GO terms and KEGG pathways. A: GO terms of DEGs in TH vs THC. B: GO terms of DEGs in P vs PC. C: KEGG pathway analysis of DEGs in TH vs THC. D: KEGG pathway analysis of DEGs in P vs PC. TH: *T. chinense*. THC: *T. chinense* chimera. PC: *P. vulgaris* chimera. P: *P. vulgaris*.

### The mobile genes between *T. chinense* and its host P*. vulgaris*

To further explore the information exchange events of genes between *T. chinense* chimera and its host *P. vulgaris* chimera at molecular level, we combined RNA-sequencing and stepwise bioinformatic classification to identify mobile transcripts between parasite plant and its host.

Consequently, 383 unigenes are most probably of *T. chinense* origin because of their absence only in P, and they are mobile genes transferred from *T. chinense* chimera to *P. vulgaris* chimera (Fig. 55 and S7 Table). The putative function classes of these parasite→host mobile genes were carbon metabolism, terpenoid biosynthesis, plant hormone signal transduction, ABC transporters, and plant-pathogen interaction (Table 3). Similarly, 285 unigenes are mobile genes transferred from *P. vulgaris* chimera to *T. chinense* chimera (Fig. 55 and S8 Table). Among them, the pathways these host → parasite mobile genes were fatty acid, steroid biosynthesis and plant-pathogen interaction (Table 3). Moreover, 189 (TH-exclusive and P-exclusive) common unigenes were finally retrieved from THC and PC (Fig. 55 and S9 Table), and these genes might be closely related to haustoria formation.

**Table 3.**
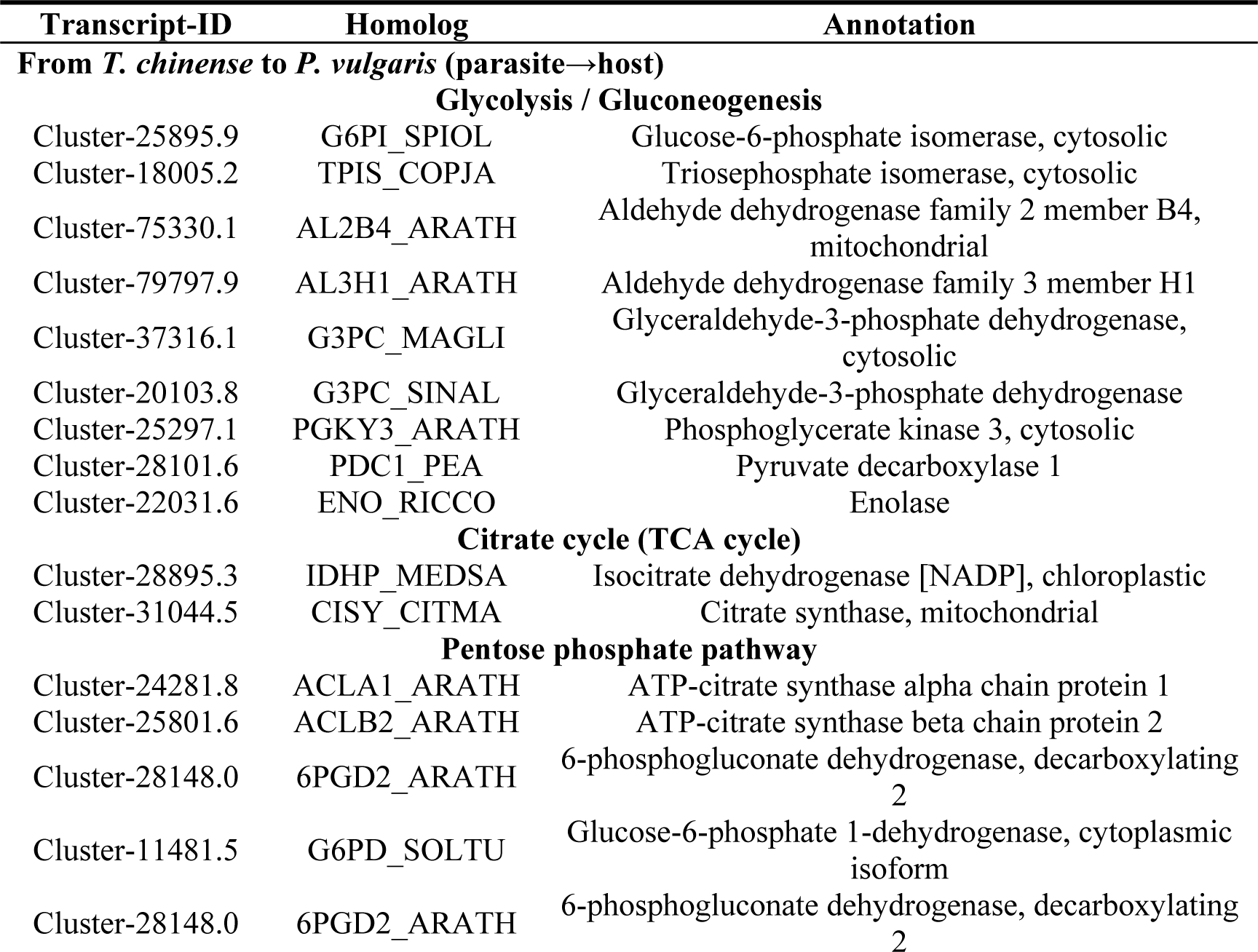
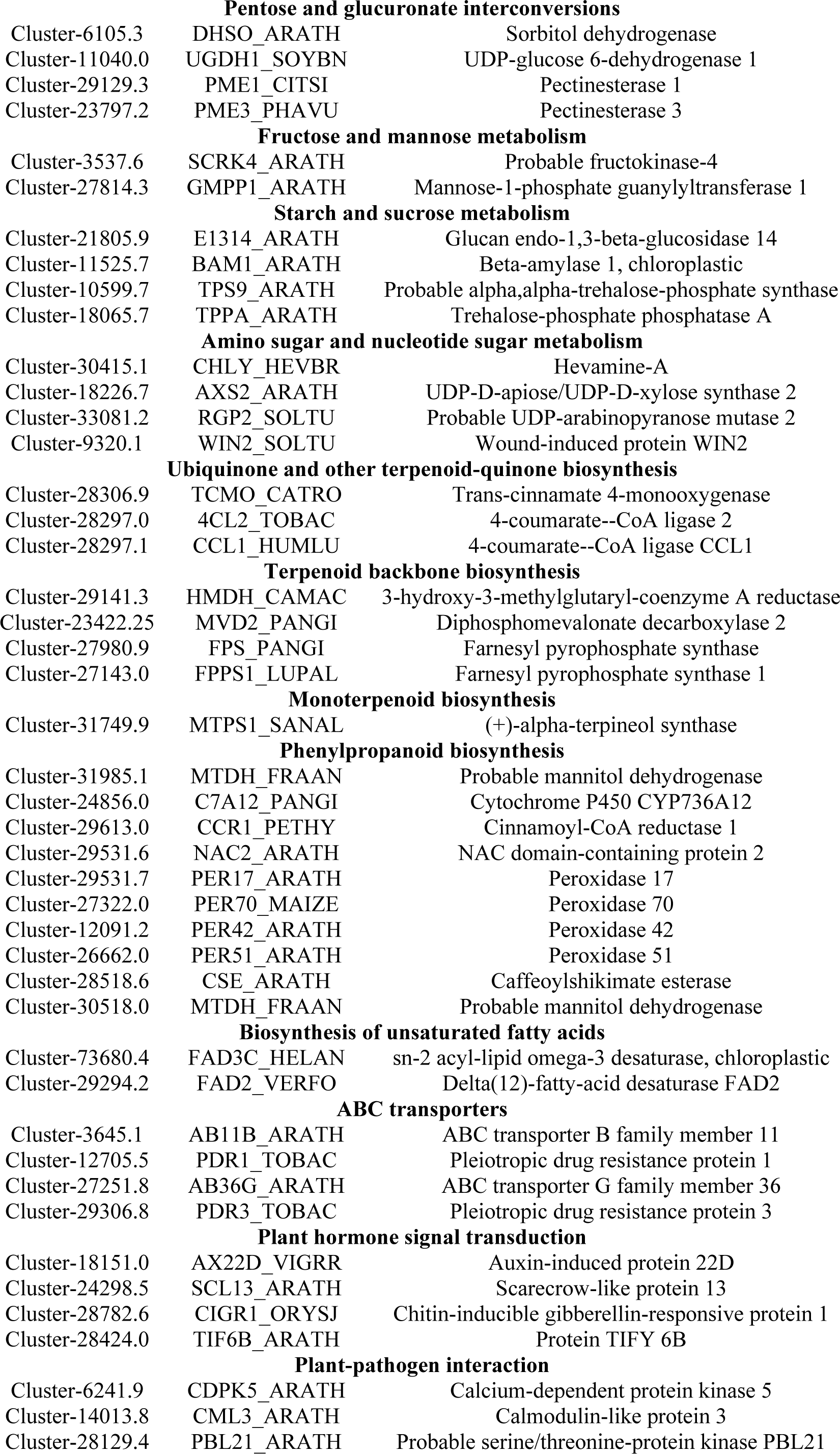
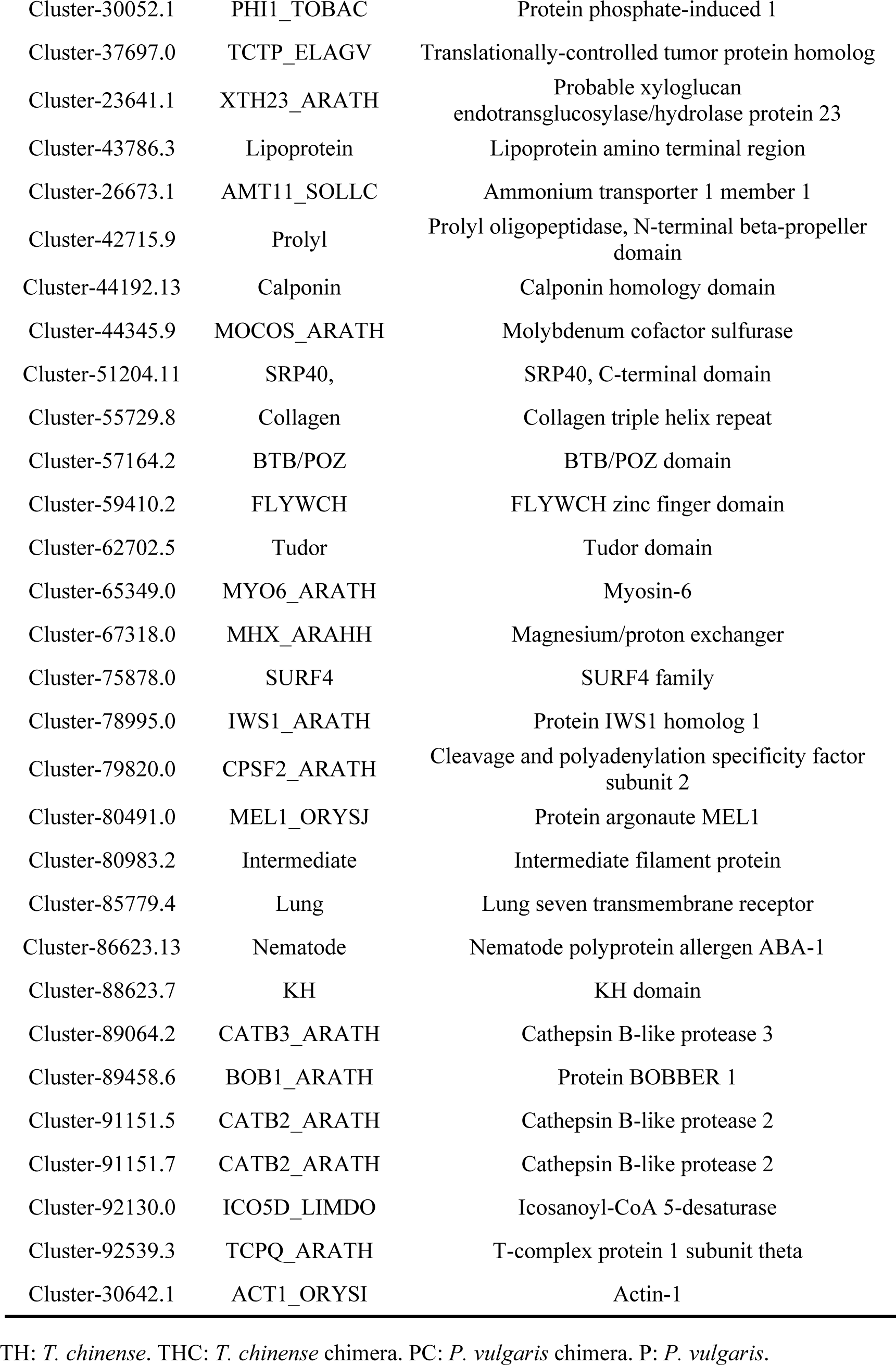
The mobile genes between *T. chinense* and *P. vulgaris*.

**Fig. 5.**
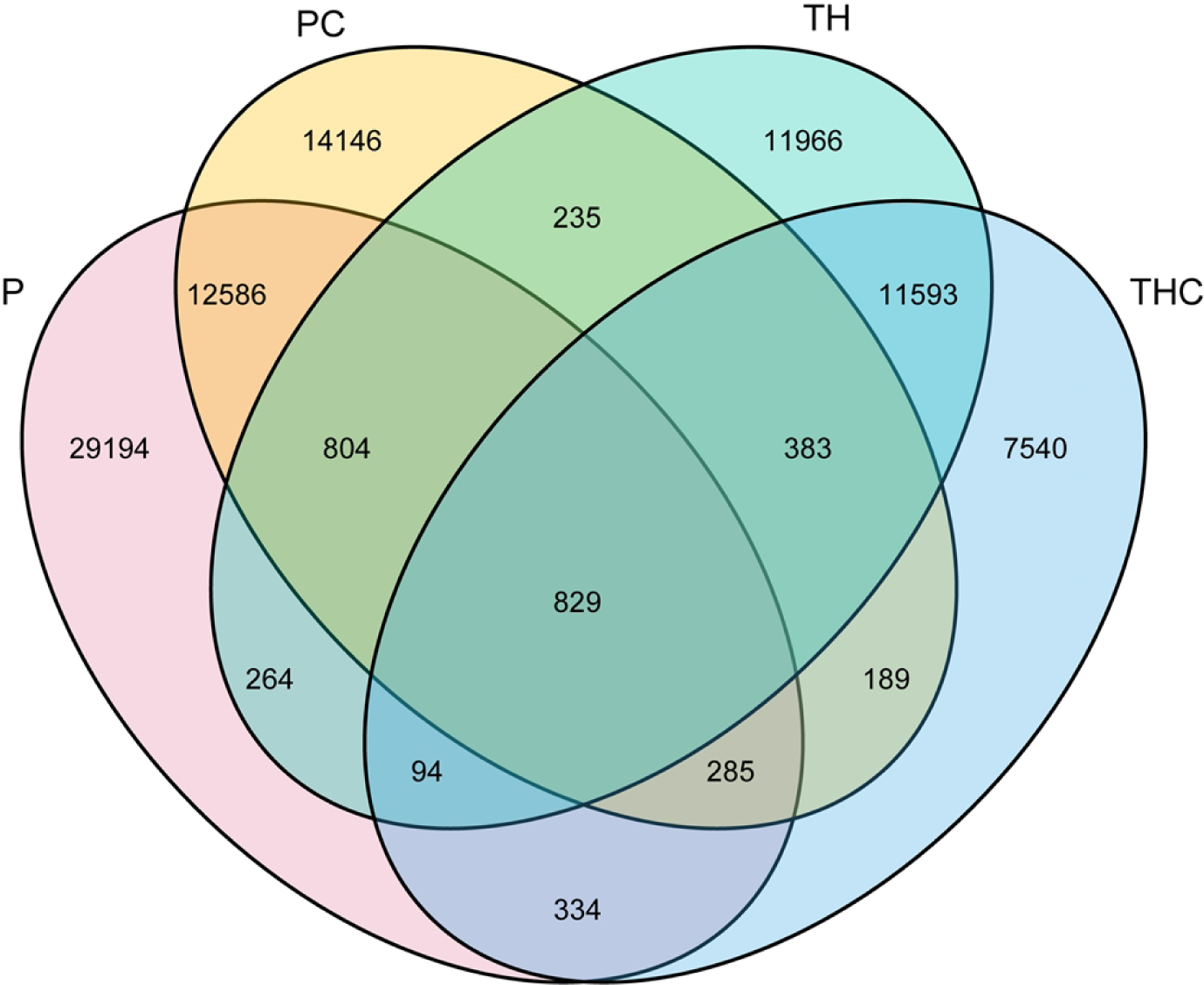
Venn diagrams showing common and unique sets of transcripts in *T. chinense* and its host *P. vulgaris* following parasition. TH: *T. chinense*. THC: *T. chinense* chimera. PC: *P. vulgaris* chimera. P: *P. vulgaris*.

### The conjoint analysis of genes and metabolites related to haustoria formation

To integrate the metabolome and transcriptome analysis, we performed canonical correlation analysis using the Pearson correlation coefficient (PCC) to show the dynamic changes in *T. chinense* and its host *P. vulgaris* post symbiosis. To systematically understand the metabolite-gene relationships ascribed to haustoria formation, we constructed the metabolite-gene network map with the threshold of |coefficient| > 0.8 (S10 Table). Out of the 189 genes linked to haustoria synthesis, 77 genes (Table 4) were carefully selected based on correlation with metabolites (DAMs related to haustoria formation). Subsequently, this narrowed down the search to 19 genes that were further analyzed for the network map.

The network map indicated that 18 genes upregulated in both THC and PC chimeras such as Peptide methionine sulfoxide reductase B5, Caffeoylshikimate esterase, UDP-glycosyltransferase 73D1, Expansin-like B1, Endochitinase and proteasome inhibitor were negatively correlated with the downregulated metabolites (Isotachioside, Tachioside, 1-O-Salicyloyl-β-D-glucose, Salicylic acid-2-O-glucoside, p-Hydroxypheny-β-D-allopyranoside, Oleic acid, Arbutin, 2,6-Dimethoxy-4-hydroxyphenol-1-O-ß-D-glucopyranoside) in both chimeras post symbiosis, and these 18 genes might be involve in haustorium formation (Fig. 6). Conversely, Protein phosphate-induced 1 downregulated in both THC and PC chimeras (Table 4) was negatively correlated with the upregulated metabolites (2,3,19-Trihydroxyurs-12-en-28-oic acid, Pinfaensic acid, 19-Hydroxyursolic acid, Jasmonic acid, 5-hydroxy-1-phenyl-7-3-heptanone) post symbiosis, and this gene might inhibit haustoria formation (Fig. 6).

**Fig. 6.**
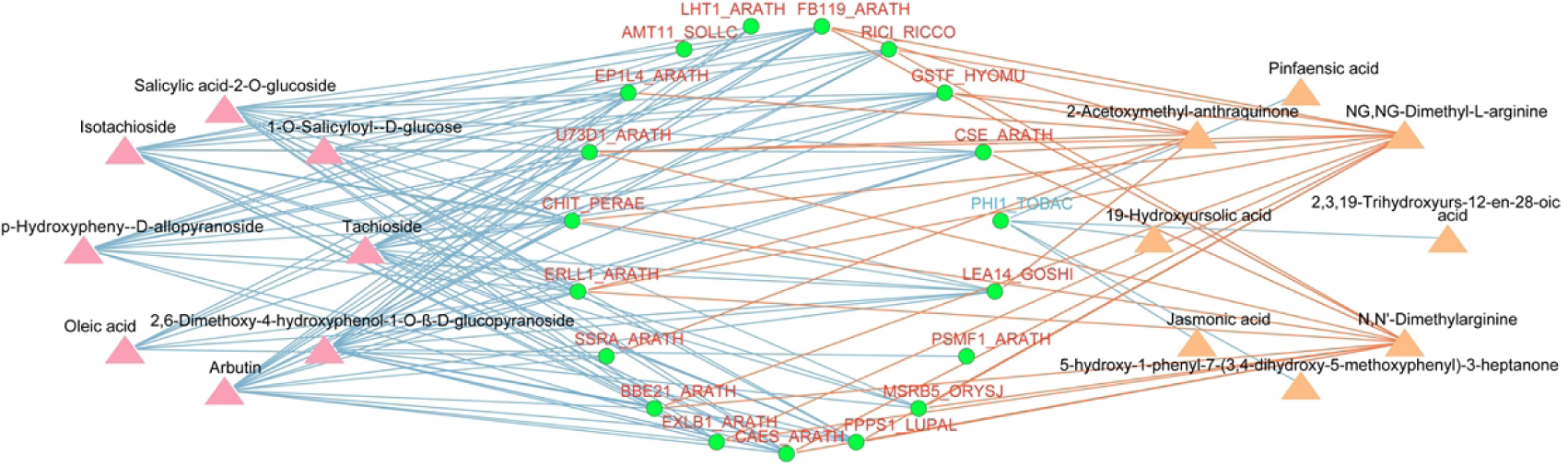
The network of metabolites and genes related to haustoria formation. Genes in the green ellipse. The upregulated genes in both THC and PC chimeras were in red font and the downregulated in blue font. Metabolites downregulated in both THC and PC are in the pink triangle and metabolites upregulated in both THC and PC are in the orange triangle. For the connection between genes and metabolites, red lines indicate the positive correlation while blue lines indicate the negative correlation. The altered pattern and annotation of haustoria formation related metabolites or genes are given in S10 Table.

**Table 4.**
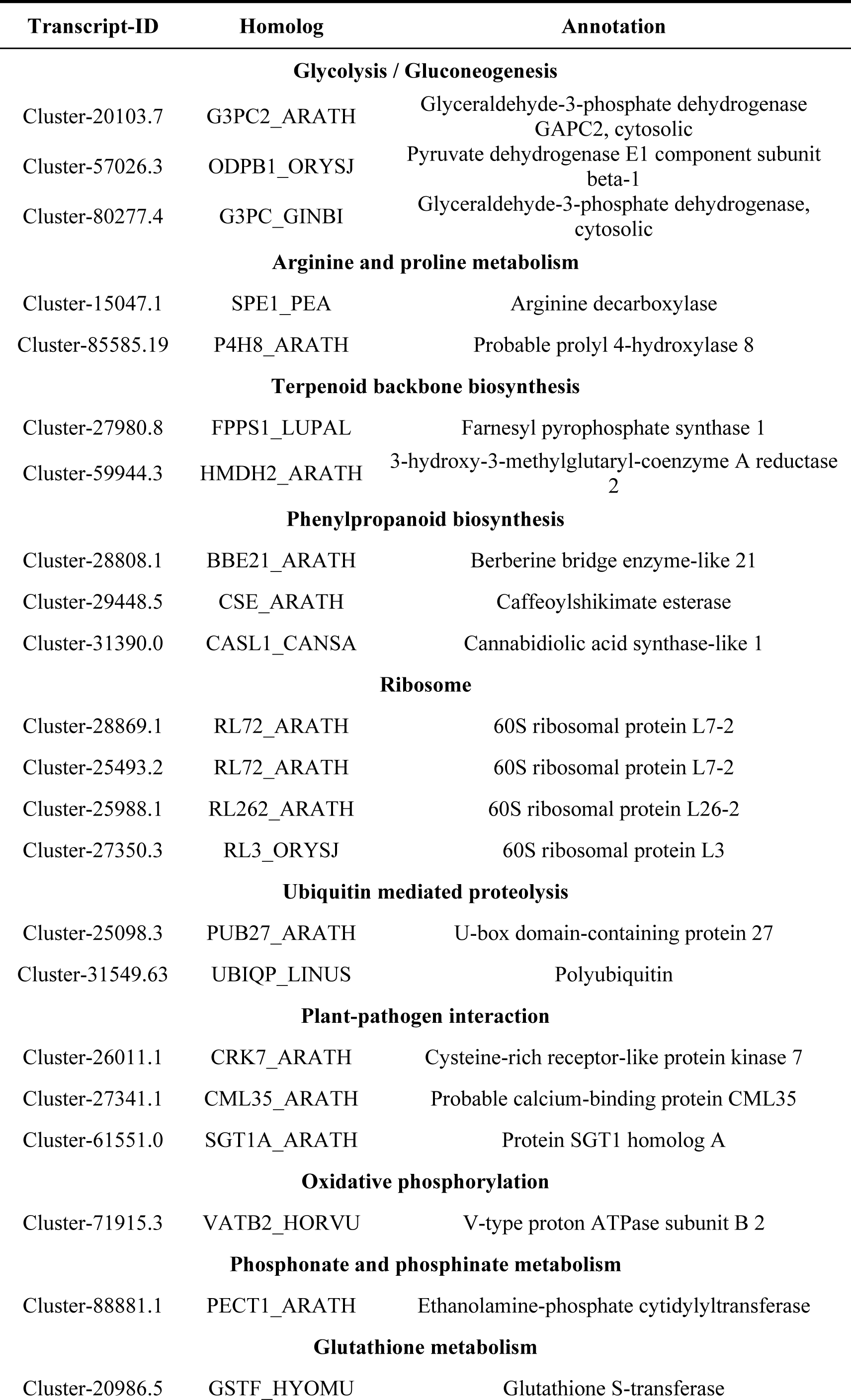
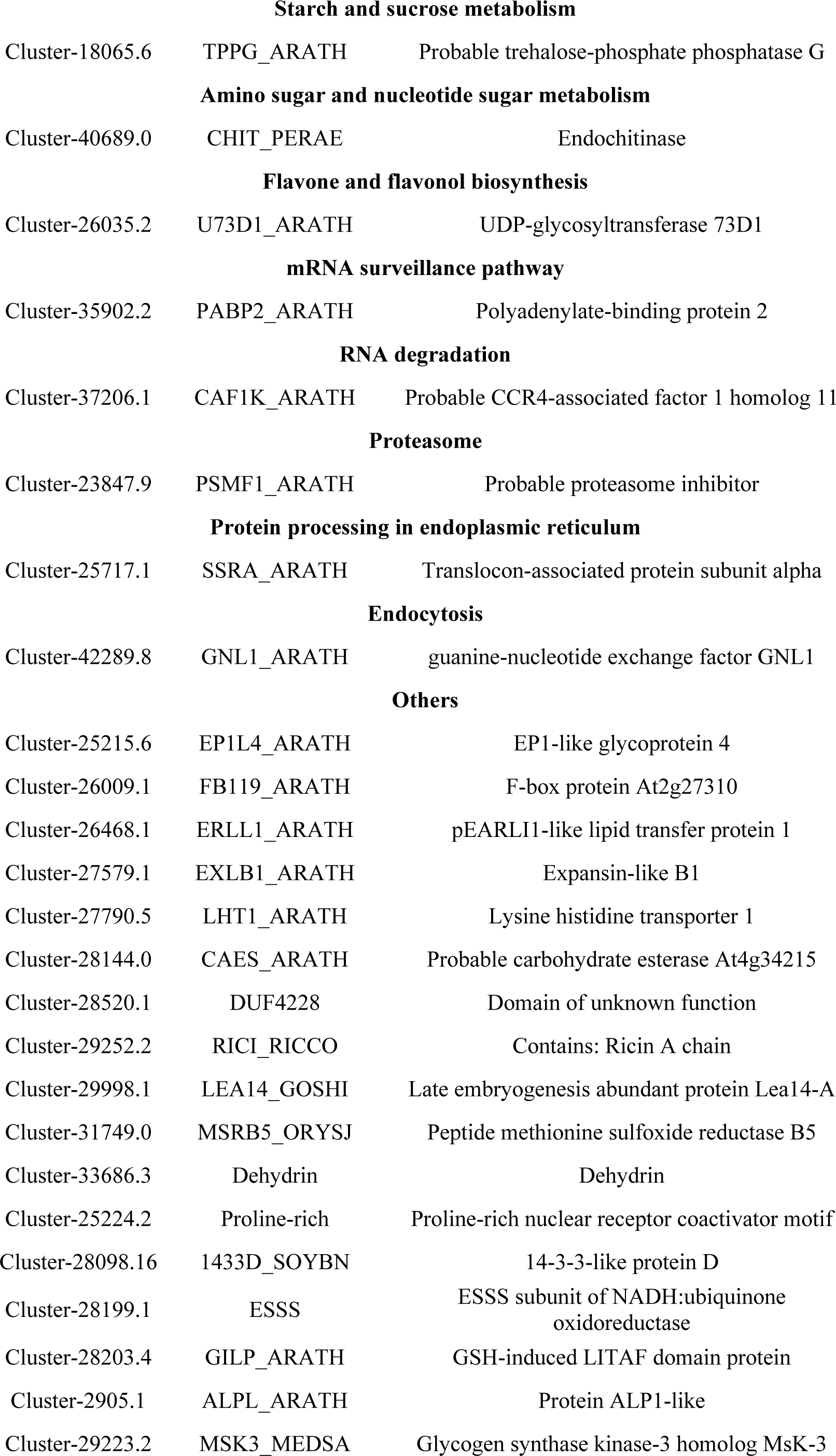
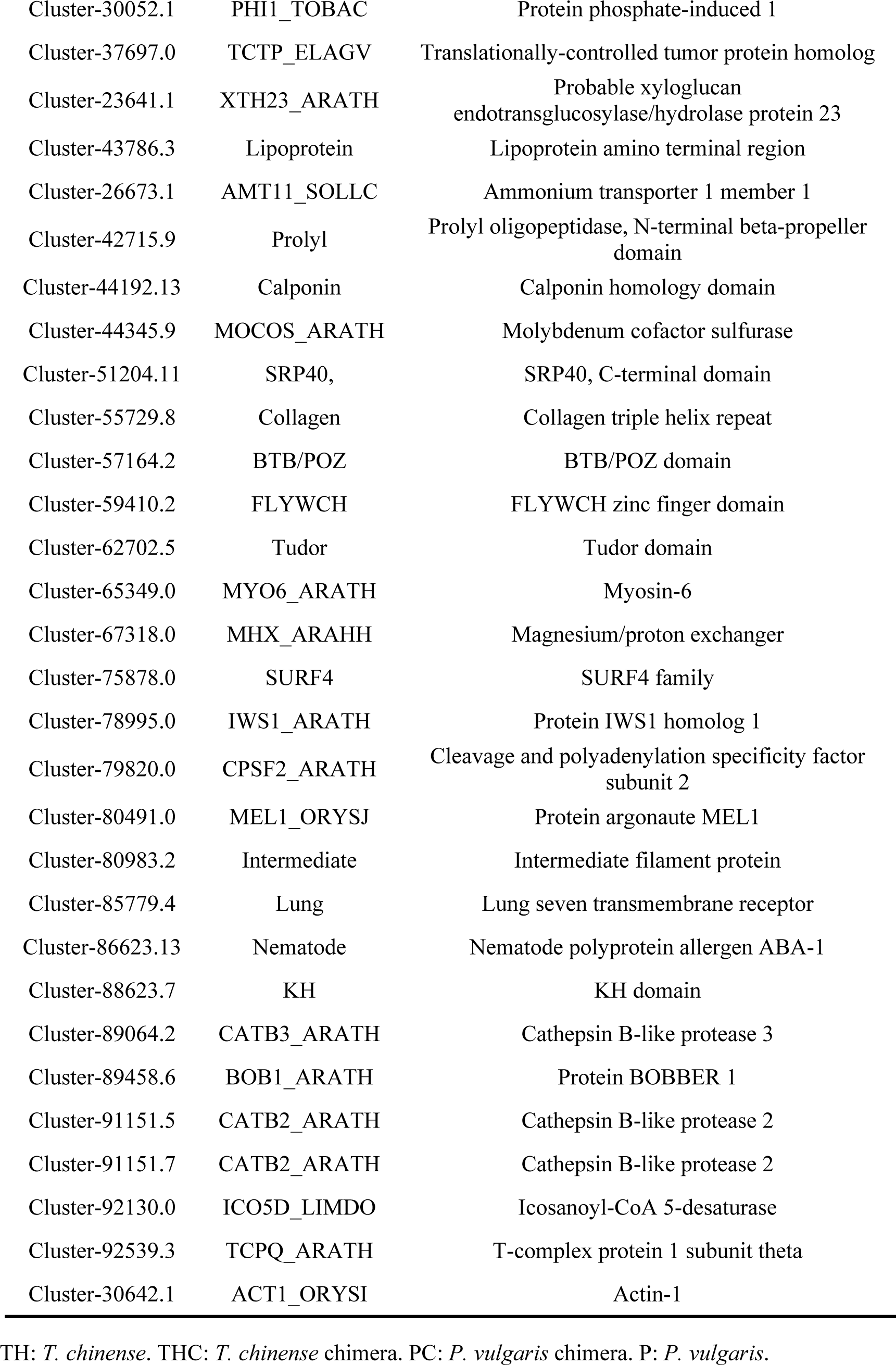
The genes correlated with haustoria formation related metabolites.

## Discussion

*T. chinense* is a medically important plant that invades its host plant through the haustoria and hijacks water, nutrients, DNA, mRNA, proteins needed to sustain its own growth and development. The essence of parasite plants’ life habits is to establish parasitic relationships with their host ^[35]^. However, to date, there has been few studies on the information exchange between *T. chinense* and its host. Therefore, this study aims to explore the changes in the metabolome and transcriptome of *T. chinense* and its host *P. vulgaris*, as well as transferred metabolites and mobile genes between *T. chinense* and its host.

According to currently available phytochemical investigations, *T. chinense* contains various compounds, with flavonoids being the main biologically active compounds responsible for its pharmacological properties and therapeutic effectiveness ^[8]^. In this study, *T. chinense* chimera showed a higher proportion of downregulated flavonoids compared with individual *T. chinense* (S2 Table). This could be a result of plant growth-defense trade-off where part of plant resources, originally allocated to growth, were redirected towards defense mechanisms, thus obtaining protective adaptation to environmental stresses. The active compounds of *P. vulgaris* mainly include flavonoids, phenolic acids, and terpenoids ^[36,37]^. After establishing a parasitic relationship, most of flavonoids and terpenoids showed an upregulation trend (S3 Table). This result indicates that parasitism promotes the accumulation of active compounds in *P. vulgaris*. These results provide a basis for understanding the metabolic mechanisms of *T. chinense*-*P. vulgaris* interactions, which will contribute to the quality control of *T. chinense*.

Phytohormones are important factors regulating plant growth and development ^[25]^. By analyzing the KEGG pathway, many DEGs in TH vs THC and P vs PC were found to be enriched in plant hormone signal transduction (Fig. 4C and 4D). Emerging studies showed that the formation of haustorium involves plant hormones such as auxin, cytokinins, and ethylene were involved in the formation of haustorium ^[38]^. Once successfully invaded, haustoria start forming xylem bridges that connect between the host and parasite xylems to initialize material transfer. Xylem bridge formation is supported by auxin flow generated by several PIN family auxin efflux carriers and AUX1/LAX influx carriers genes expressed within invading haustoria ^[39]^. Haustorium-inducing factors (HIFs) trigger the expression of an auxin biosynthesis gene in root epidermal cells at the haustoria formation sites. This process leads to cell division and expansion, resulting in the formation of a semi-spherical pre- or early haustorium structure ^[40]^. Therefore, the auxin biosynthesis/signaling-related genes were highly abundant in *T. chinense* haustoria ^[34]^. Furthermore, the auxin response could be one of the shared mechanisms in haustoria formation among parasitic plants ^[41]^. In the present study, the levels of auxins such as indole, 3-indolepropionic acid and 3-indoleacrylic acid decreased in *T. chinense* chimera (Table 1). Similarly, a downtrend of auxin content was detected in *Cuscuta japonica* during its parasitization ^[25]^. The auxin pathway may play an important role in the host-parasite association ^[41]^. Therefore, auxin transport may participate in establishing the host-parasite association. JA is an ancient regulator that controls the biosynthesis and/or transport of systemic signals, and it plays a crucial role in the biosynthesis or transport of between-plant mobile signals ^[25]^. Furthermore, the host JA signaling plays a role in regulating the gene expression in the parasitizing *Cuscuta* ^[41]^. In this study, JA was upregulated in THC and PC (Table 1). However, the specific functions of JA in parasitic plants remain unexplored. We speculate that the increased JA levels in *T. chinense* chimera may be related to its defense mechanism against the host, as the chimera also accumulating more JA. In short, the progression of haustorium organogenesis and the host-parasite interaction is controlled by phytohormones. To understand how plants coordinate multiple hormonal components in response to diverse developmental and environmental cues represents a significant challenge for the future. In our study, the metabolic changes caused by *T. chinense* parasitism were associated with phenylpropanoid biosynthesis, flavonoid biosynthesis, plant hormone signal transduction, fructose and mannose metabolism, vitamin B6 metabolism and photosynthesis-antenna proteins (Fig. 4C). The fructose and mannose metabolism pathway is crucial for the success of parasitism ^[27]^. In the case of *Orobanche aegyptiaca*, the host-induced suppression of the mannose 6-phosphate reductase gene is concomitant with significant mannitol decrease and increased tubercle mortality ^[42]^. In plant-pathogen interaction, the pathogen secretes mannitol as a buffer against oxidative stress, and the host plant activates mannitol dehydrogenase to counter it ^[43]^. In the study, the relatively high mannitol level in *P. vulgaris* chimera might be a consequence of this host-parasite interaction (S3 Table).

Parasitic plants and their hosts are often phylogenetically very distant, and the haustoria establish physical and physiological connections between the host and parasitic plants, thereby dominating most of their interactions ^[6]^, making the host-parasite systems very suitable for the identification of mobile substances. Secondary metabolites are essential for plant survival and are typically biosynthesized in specific tissues and cell types before being transported to neighboring cells or even to other tissues or other organs ^[44]^. Some secondary metabolites in the host can be transferred to the parasite plant ^[25]^. We have identified 4 metabolites that were transferred from *P. vulgaris* chimera to *T. chinense* chimera (Table 2). In this study, 2-ethylpyrazine was identified to be the transferred metabolite from *T. chinense* chimera to *P. vulgaris* chimera (Table 2). Although what effect 2-ethylpyrazine has on the parasitism relationship remains unknown, we speculate that it may be a metabolite of *T. chinense* that attracts host plants and successfully colonizes them. Actually, how parasitic plants accept secondary metabolites from their hosts and the ecological impact of the translocated secondary metabolites in parasitic plants require further exploration ^[25]^. In future experiments, we can apply 2-ethylpyrazine to *P. vulgaris* or other host plants of *T. chinense* and observe whether *T. chinense* can colonize faster or promote its growth to verify the role of 2-ethylpyrazine in contributing to establish parasitism relationship.

Compared to other host-pathogen systems ^[45]^, there were few reports on the interactions between parasite plant-host ^[46]^. An important aspect of the host-parasite interaction is the effect of host’s growth stage and environment on the mobile mRNAs expression ^[23]^. The presence of haustoria also facilitates the transfer of RNAs between parasitic plants and the host ^[47]^. RNA-sequencing analysis has indicated the trafficking of thousands of mRNA species between hosts and *Cuscuta pentagona* ^[48]^. There was also large-scale mobile mRNA between *Haloxylon ammodendron* and the parasitic plant *Cistanche deserticola* ^[46]^, and the mRNA abundance gradient is likely to be the determinant of mobility ^[25]^. In this study, cross-species mRNA movement was identified between *T. chinense* and *P. vulgaris*, and 383 and 285 mobile mRNAs might be transferred from *T. chinense* and *P. vulgaris* to host and parasite through haustoria, respectively (S7 and S8 Table). However, these mobile transcripts need further investigation.

## Conclusion

The study presents a comprehensive analysis of the metabolome and transcriptome of *T. chinense*, *P. vulgaris* and their chimeras. The identification of 5 transferred metabolites and 668 mobile genes exchanged between *T. chinense* and P*. vulgaris* underscores the complexity of the parasitic relationship, revealing a substantial inter-organismal transfer of resources and genetic information. The discovery of 56 metabolites and 189 genes related to haustoria formation further emphasizes the intricate biological processes underpinning the establishment of parasitism. The constructed gene regulatory network has pinpointed 18 genes that promote and one gene that inhibit haustoria formation, highlighting potential targets for further investigation into the mechanisms of parasitic plant development. The critical role of the fructose and mannose metabolism pathway in parasitism success is also emphasized. These findings indicate a strategic exploitation of host resources that are likely essential for the parasitic plant’s survival and proliferation.

In conclusion, our results suggest that *T. chinense* engages in a sophisticated and dynamic biological exchange with *P. vulgaris*, utilizing both metabolites and mobile mRNAs to facilitate haustoria formation and successful parasitism. The elucidation of these complex interactions not only expands our understanding of the molecular dialogues between parasitic and host plants but also sets the stage for future explorations into controlling or harnessing these interactions for agricultural and ecological benefit.

## Materials and methods

### Plant materials and sample collection

Five plants each of *T. chinense*, *P. vulgaris* and their commensal chimera were randomly selected for sampling independent roots and chimeric roots, and the *T. chinense* chimera (THC) and *P. vulgaris* chimera (PC) were sampled from the symbiont roots post parasitization. To minimize any surface tissue contamination, the sampled roots or chimera with three biological replicates were washed 1-2 times with PBS/RNase-free water, and frozen in liquid nitrogen and stored at −80°C for the subsequent metabolomic, transcriptomic analysis.

### Metabolome analysis

For the widely targeted metabolomic profiling, four type of root samples aforementioned were freeze-dried by vacuum freeze-dryer (Scientz-100F). The freeze-dried sample was crushed using a mixer mill (MM 400, Retsch) with a zirconia bead for 1.5 min at 30 Hz. Dissolve 50 mg of lyophilized powder with 1.2 mL 70% methanol solution, vortex 30 seconds every 30 minutes for 6 times in total. Following centrifugation at 12000 rpm for 3 min, the extracts were filtrated (SCAA-104, 0.22 μm pore size; ANPEL, Shanghai, China, http://www.anpel.com.cn/), then analyzed using an UPLC-ESI-MS/MS system (UPLC, ExionLC™ AD, https://sciex.com.cn/; MS, Applied Biosystems 6500 Q TRAP, https://sciex.com.cn/) ^[49,50]^.

Based on the mass spectrometry data, metabolites were identified using the Metware Database (MWDB, Wuhan, China) (www.metware.cn) and quantified according to peak intensity. Both unsupervised principal component analysis (PCA) and orthogonal projections to latent structure-discriminant analysis (OPLS-DA) were used to observe the overall differences in metabolic profiles between groups to identify their significant differential metabolites. The quantification data of metabolites were normalized by unit variance scaling and used for the subsequent analysis (http://www.r-project.org/) ^[51]^.

### Screening of differentially accumulated metabolites

To determine the metabolomic differences of *T. chinense* and its host post parasitization, the differentially accumulated metabolites (DAMs) in the TH vs THC and P vs PC groups were screened. Variable importance in projection (VIP) values were extracted from OPLS-DA results, those selected and metabolites with VIP ≥ 1 and absolute |log_2_FoldChange| ≥ 1 were defined as DAMs ^[52]^.

The DAMs were annotated using the KEGG Compound database (http://www.kegg.jp/kegg/compound/) and mapped to the KEGG Pathway database (http://www.kegg.jp/kegg/pathway.html) ^[53]^. Then a KEGG pathway enrichment analysis was performed, and the significance was determined by hypergeometric test p-values ≤ 0.05.

### RNA extraction, library construction, and sequencing

Total RNA was isolated using the Trizol Reagent ^[54]^ (Invitrogen Life Technologies, Shanghai, China). To ensure the RNA samples were integrated and DNA-free, agarose gelelectrophoresis was performed. RNA purity was then determined by a nanophotometer. Following that, a Qubit 2.0 Fluorometer and an Agilent 2100 BioAnalyzer were used to accurately measure RNA concentration and integrity, respectively. The qualified samples were processed with oligo (dT) beads to enrich the mRNA, which was broken into fragments and used as templates for the cDNA library. To qualify the cDNA library, the fluorometer was used for primary quantification and the bioanalyzer was then used to insert text size. The qualified library was sequenced using the Illumina HiSeq 6000 platform.

### RNA-Seq analysis

Clean reads were obtained by eliminating low-quality reads and assembled using Trinity 2.8.5 software ^[55]^. The transcripts were assembled and then clustered into unigenes. The method of fragments per kilobase of transcript per million fragments mapped (FPKM) was applied to calculate the expression levels of genes. DESeq2 ^[56]^ was used to identify differential expression genes (DEGs) based on the thresholds of the adjusted p-value padj < 0.05 and |log_2_FoldChange| ≥ 1 ^[56]^. Then DEGs were annotated by the NR, SwissProt, GO, KOG, Pfam, and KEGG databases. Finally, GO and KEGG pathway enrichment analysis were performed on DEGs to reveal functional modules and signal pathways of interest.

### Integrated metabolomic and transcriptomic analysis

Based on the metabolite content and gene expression data, Pearson correlation tests were used to detect associations between gene expression and metabolite content. Correlations between DAMs and DEGs were filtered according to Pearson correlation coefficient (PCC) and *P*-value. Only the significant associations with |PCC| > 0.80 and *P*-value < 0.05 were selected for constructing network of metabolome and transcriptome, and the metabolite-gene relationships related to haustoria formation were visualized using Cytoscape (v3.9.0) ^[57]^.

## Supporting information

S1 Table. Metabolites in the *T. chinense*, *P. vulgaris* and their chimeras.

S2 Table. The differentially accumulated metabolites (DAMs) in the *T. chinense* and *T. chinense* chimera.

S3 Table. The differentially accumulated metabolites (DAMs) in the *P. vulgaris* and *P. vulgaris* chimera.

S4 Table. The differentially accumulated metabolites (DAMs) related to haustoria formation.

S5 Table. The differentially expressed genes (DEGs) in the *T. chinense* and *T. chinense* chimera.

S6 Table. The differentially expressed genes (DEGs) in the *P. vulgaris* and *P. vulgaris* chimera.

S7 Table. Mobile 383 genes transferred from *T. chinense* chimera to *P. vulgaris* chimera.

S8 Table. Mobile 285 genes transferred from *P. vulgaris* chimera to *T. chinense* chimera.

S9 Table. The 189 genes related to haustoria formation.

S10 Table. The correlation of haustoria formation related genes and metabolites.

## Acknowledgments

We would like to express our gratitude to the members of our laboratory for their suggestions and guidance on the manuscript, and to Metware Biotechnology Co., Ltd. (Wuhan, China) for essential technical support.

## Author Contributions

Conceptualization: Mingpu Tan, Zengxu Xiang.

Data curation: Anping Ding, Mingpu Tan, Zengxu Xiang.

Investigation: Anping Ding, Ruifeng Wang.

Methodology: Anping Ding, Ruifeng Wang, Juan Liu, Wenna Meng.

Resources: Anping Ding, Ruifeng Wang, Zengxu Xiang.

Software: Anping Ding, Ruifeng Wang, Juan Liu, Wenna Meng.

Validation: Anping Ding, Ruifeng Wang.

Visualization: Anping Ding, Juan Liu, Wenna Meng, Gang Hu.

Writing – original draft: Anping Ding, Mingpu Tan.

Writing – review & editing: Mingpu Tan, Zengxu Xiang.

All authors have read and agreed to the published version of the manuscript.

## Data availability

The RNA-seq raw data used in this study have been deposited in Sequence Read Archive (SRA) database in NCBI under accession number PRJNA1054813 (https://www.ncbi.nlm.nih.gov/sra/PRJNA1054813, which will be released upon publication).

